# Amnesia Induced by Suppressing the Focus of Internal Attention

**DOI:** 10.64898/2026.11.03.763713

**Authors:** Akul Satish, Justin C. Hulbert, Michael C. Anderson

## Abstract

Our thoughts can recede as we direct attention to the external world. This experience may reflect the suppression of thoughts about past events, future worries, or stray ideas that could otherwise disrupt focus. Such thoughts may arise spontaneously from replay mechanisms supported by the episodic memory system; hence, sustaining external focus may require suppressing these systems to stop internal distraction. Here, periods of high external focus disrupted diverse episodic content activated near in time, inducing an ‘amnesic shadow’. This shadow occurred for both novel content and already formed memories and exhibited key markers of active suppression: forgetting beyond expectations from time alone, which generalised to an alternative reminder. External attention therefore automatically engages an active suppression mechanism that appears to broadly inhibit cognitive systems supporting the emergence of distracting thoughts. The findings raise the possibility that people may naturally gravitate toward activities demanding external attention, unwittingly, to regulate unwanted thoughts.

## Introduction

Directing attention to the external world may require suppressing our thoughts. When we focus on the outside world, such as observing the colours of an evening sky or listening to birdsong, our stream of thoughts and worries can recede. What, then, is the fate of our thoughts when attention shifts outward? Do they simply pause, remaining ready to be re-engaged, or are they actively suppressed?

Suggestive evidence comes from the brain’s default network, which supports internally directed cognition involved in generating thought content such as memories, future simulations, self-relevant thoughts, and worries^1–6^. The default network exhibits increased activity in situations that demand access to internal content, regardless of whether attention is directed inward^4,7^ (e.g., self-reflection) or outward^2,8–10^ (e.g., memory-guided navigation). However, in situations with minimal requirement for internal content, such as maintaining attention towards external sensory input, default network activity is reduced^11–17^, consistent with the possibility that external sensory attention suppresses mentation. Yet, such activity reductions could also occur if internal mentation is simply paused and not actively suppressed.

To illustrate these alternative possibilities, consider the cognitive mechanisms that could determine the state of internal mentation during external focus^18,19^. Thought contents may be preserved if mentation pauses when internal attention is withdrawn and redirected externally, which we refer to as the attentional withdrawal hypothesis. Alternatively, thought contents may be disrupted if internal mentation is inhibited to support external focus, which we refer to as the active suppression hypothesis (see Figure 1).

**Figure 1.**
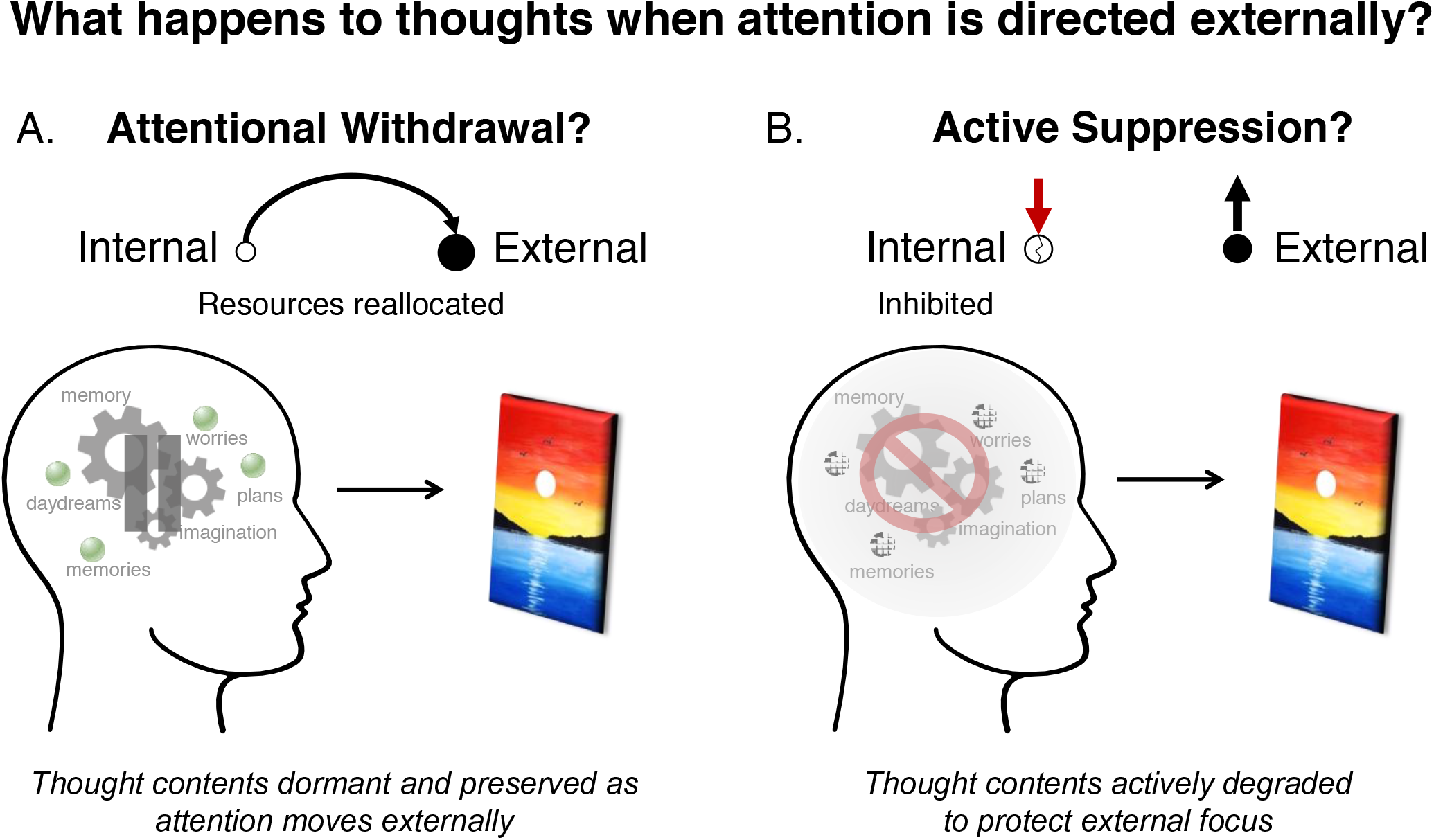
Two mechanisms that may determine the state of thoughts during external sensory attention. Internally directed cognition is represented by processes such as memory and imagination (gears) that activate thought contents (orbs) such as memories, worries, plans, and daydreams. (A) Under the attentional withdrawal account, internal attention is withdrawn and reallocated to external perceptual input (sunset), leaving thought contents dormant and ready to be re-engaged. (B) Under the active suppression account, internally directed processes are actively inhibited, degrading thought contents and protecting external focus (sunset).

Whether internal mentation is paused or suppressed may depend on whether the persisting accessibility of thoughts under-mines external attention. Indeed, there is ample evidence suggesting that external task performance is worse when thought contents intrude and involuntarily redirect attention inwardly. Vivid examples of this competition come from situations where internal contents arise spontaneously and uninvited^20^ during mind-wandering^7,21^, reflected by increased default network activity^22,23^ and measurable costs for external attention. Greater mind-wandering is associated with worse performance on sustained attention tasks and reduced neural processing of external stimuli^24–27^. These costs are further amplified in psychiatric disorders characterised by intrusive thinking and rumination, where default network hyperactivity is a consistent finding alongside poor external task performance^28–31^. Together, these observations suggest that internal contents can interfere with external attention^32,33^, and maintaining external focus may require limiting their interference.

How does internal content interfere with external attention? One possible mechanism is attentional capture, whereby salient or unexpected stimuli generate a priority signal that reflexively reorients attention to that stimulus and disrupts ongoing task performance^34–39^. An abrupt flash of light, for instance, will pull your attention away from reading this article via attentional capture mechanisms. Whereas attentional capture is well established for external stimuli, recent evidence suggests that internal content such as episodic memories can engage the same reflexive mechanisms to involuntarily intrude into awareness^40–42^. In doing so, episodic memories may compete with external sensory attention for shared attentional resources^8,43–50^ and disrupt external focus.

Evidence from attentional capture has important implications for clarifying the attentional withdrawal versus suppression hypotheses. When attention is focused externally, episodic memories could involuntarily pull attention back inward^40,42^, suggesting that merely withdrawing internal attention would leave thought contents such as episodic memories free to recapture attention. An active suppression mechanism may therefore be necessary to inhibit internally directed processing, thereby degrading the accessibility of thoughts and stopping them from capturing attention and disrupting external focus.

Diverse thought contents arise from activity in the episodic memory system, often in the absence of obvious cues or deliberate intention^20,40,51–53^. Specifically, internal content may spontaneously arise via hippocampal replay^54–60^, which supports the recombination of previously encoded experiences into varied thought content. Inhibiting the episodic memory system would stop replay mechanisms, preventing internal content from becoming active and drawing attention. Consistent with this possibility, the hippocampus, part of the wider default network, exhibits reduced activity when attention is focused on sensory input^3,61^. Hence, episodic memory processes may be suppressed during external attention.

If external attention broadly suppresses episodic memory processing, this suppression should have measurable consequences that extend beyond the period of external focus itself. For instance, episodic content activated before periods of external focus may be poorly retained if hippocampal stabilisation processes are disrupted. Similarly, episodic content encountered after periods of external focus may be poorly encoded if hippocampal encoding processes are suppressed. Hence, measuring retention for episodic content activated either before or after periods of external focus offers a behavioural assay of whether episodic memory processing is suppressed during external attention. External attention may therefore induce an ‘amnesic shadow’ for thought contents arising before or after periods of demanding external focus.

Precedent for an amnesic shadow exists in paradigms that ask participants to intentionally stop retrieval of an unwanted memory, in response to a reminder. Such retrieval stopping^40,62,63^ reduces retention for entirely unrelated ‘bystander’ content encountered both before and after retrieval stopping events^64–68^. Because bystanders in these studies are unrelated to the suppressed memories, memory deficits are unlikely to reflect targeted suppression of the memory instigating retrieval stopping, but instead suggest a global suppression of episodic memory processing. Supporting this process inhibition, the magnitude of the amnesic shadow increased with greater hippocampal downregulation during retrieval stopping^65^. Hence, episodic memory processing can be suppressed by retrieval stopping, providing a paradigm to test for an amnesic shadow due to sensory attention.

In the current study, we adapted this paradigm to test whether episodic memory processing is suppressed during external attention by replacing the retrieval stopping task with a visual discrimination task demanding high sensory attention (see Figure 2). Specifically, participants completed a difficult visual discrimination task in which they identified the brightest of four closely shaded boxes (adapted from Law et al.^61^). Detecting the target box demanded sustained attention on boxbrightness with minimal reliance on internal content (see Figure 2A). In an easy version, the target was clearly brightest, placing low demands on external sensory attention and serving as a control condition. Across three experiments, episodic content was presented both before and after difficult discriminations, serving as ‘bystanders’ to periods of hypothesised episodic memory suppression. If memory processes are suppressed during external attention, bystanders to difficult discriminations should be subsequently remembered more poorly than bystanders to easy discriminations (see Figure 2B).

**Figure 2.**
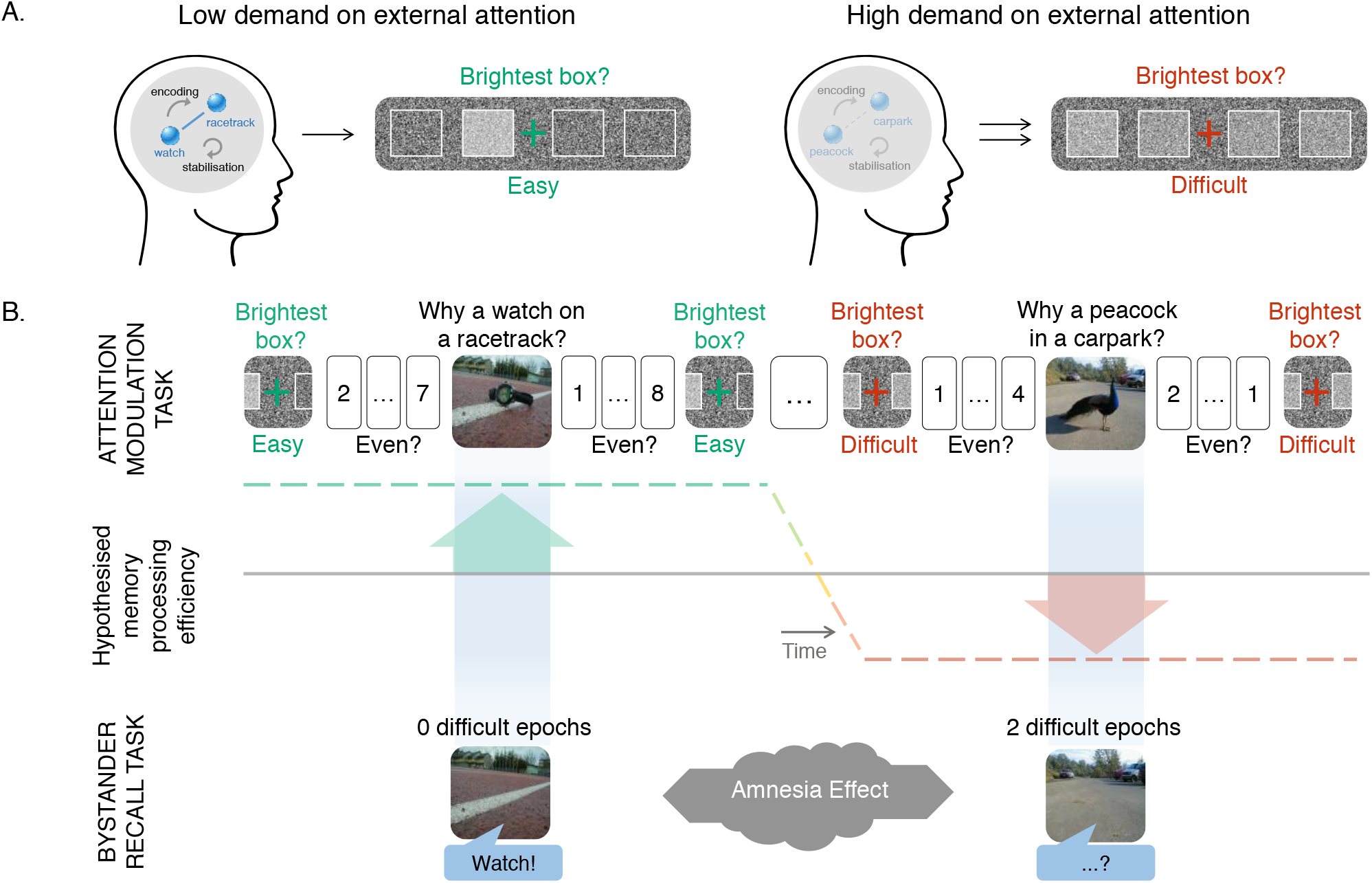
(A) Schematic illustration of the visual discrimination task under low and high demand on external attention. When demand is low (easy discrimination), memory processes such as encoding and stabilisation may remain active (left). When demand is high (difficult discrimination), memory processing may be reduced (right). (B) Phase 1 (attention modulation phase) includes discrimination trials interleaved with novel ‘bystander’ object-scene images that participants encode by silently answering the question, “Why is the pictured object in this location?”. Odd/even buffer trials completed before and after each bystander judgement match the task switching costs across easy and difficult discriminations. The dashed-line graph illustrates the predicted efficacy of memory processing during the attention modulation phase. In phase 2 (bystander recall), participants attempt to recall the bystander objects when cued by the scenes; the key prediction is that bystanders surrounded by two difficult discrimination epochs are recalled worse than those surrounded by zero difficult discrimination epochs.

The first experiment tested whether the formation of new episodic memories was impaired for events presented close in time to difficult relative to easy discriminations. The second experiment examined whether such impairments were specific to the episodic context in which bystanders were learnt^65,69^ relative to memory for the item itself. Such findings would suggest that focusing externally has behavioural consequences for nearby episodic content, providing evidence for an amnesic shadow and demonstrating modulation of episodic memory processing.

The third experiment was specifically designed to distinguish between attentional withdrawal and active suppression (adapted from Zhu et al.^67,68^). Previously formed memories were reactivated near in time to difficult (and easy) discriminations. Subsequently, their retention was compared against baseline memories that were initially learned but not reactivated between discriminations at all. If attention was simply withdrawn during difficult discriminations, bystander memories would remain intact and their retention should resemble that of baseline memories. If episodic memory processing was actively suppressed during difficult discriminations, retention of bystander memories should fall below that of baseline memories^63^. Furthermore, prior work suggests that suppressed memories remain inaccessible even when probed with an independent cue^68,70^, a property known as cue-independence. To test for cue independence, bystander memories were initially learned with an alternative cue, distinct from the cue used to reactivate bystanders. Observing below-baseline forgetting and cue independence would provide evidence for the degradation of episodic memories when attention is focused externally, indicating the involvement of suppression. Thus, if nearby episodic memories are degraded when attention is focused externally, then episodic memory processing, a key form of internal mentation, is actively suppressed rather than merely neglected.

## Results

### Impaired formation of episodic memories

Experiment 1 implemented a difficult visual discrimination task to encourage external focus, which we hypothesised would induce an amnesic shadow, relative to an easy discrimination task that had a low demand on external attention. Both easy and difficult visual discriminations were presented in the attention modulation phase, interleaved with bystander images consisting of uncommon objects embedded in background scenes (e.g., a peacock in a carpark). Participants covertly generated a story explaining how the object (a peacock) came to appear in that scene (a carpark), which encouraged object–scene associative memory formation, before they rated their difficulty in generating these stories. In a subsequent surprise memory test of bystanders, only the background scenes were shown, and participants recalled the associated object (see Figure 2).

If difficult discriminations hinder the retention of bystander content, their recall should be worse than bystanders near easy discriminations. In line with this prediction, bystander recall was worse when surrounded by two relative to zero difficult discrimination epochs: *F*(1, 96) = 20.57, *p* < .001, *η*_*p*_^*2*^ = .18, 90% CI = [.08, .29] (see Figure 3).

**Figure 3.**
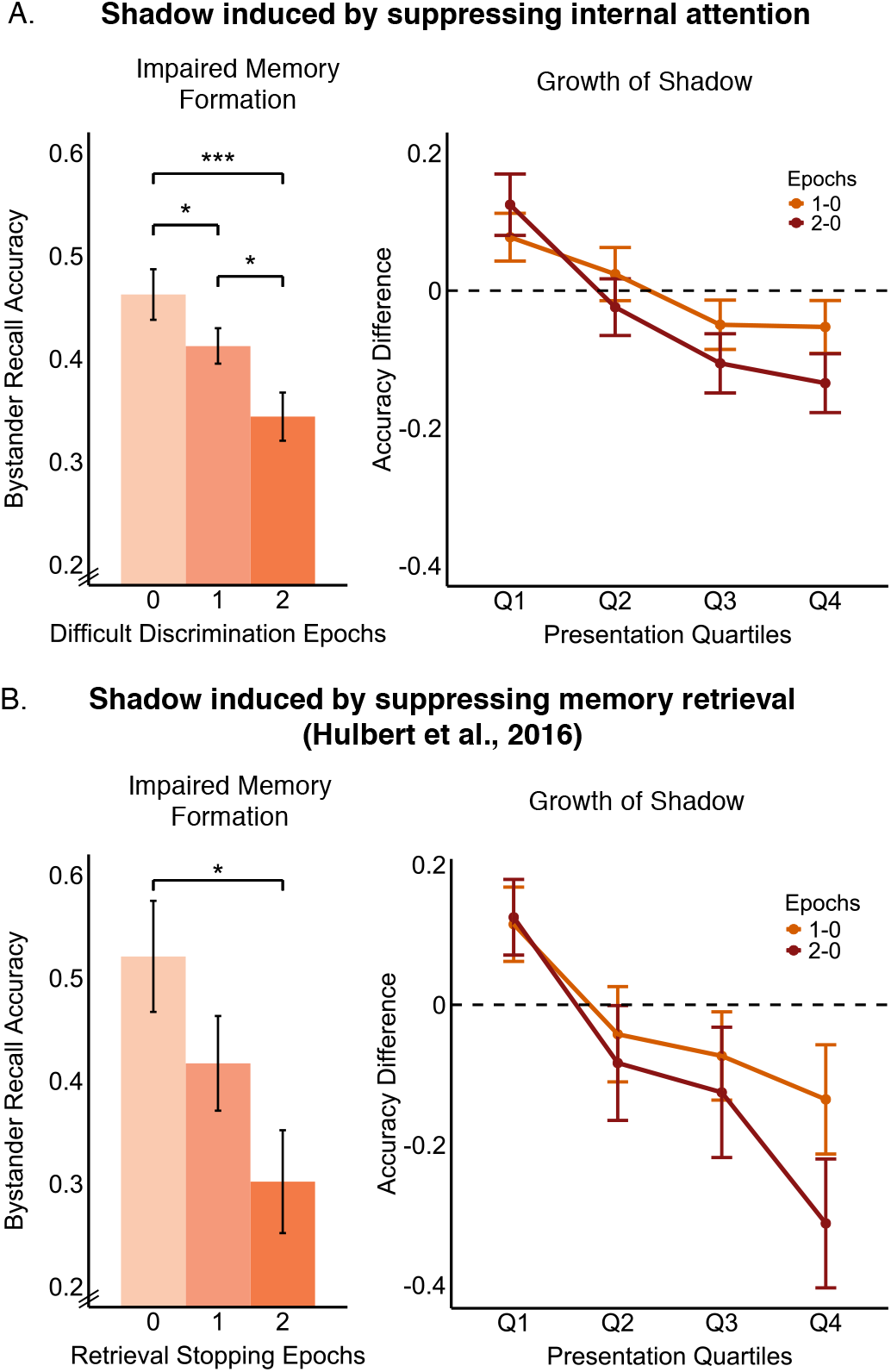
(A) The amnesic shadow for episodic memory formation induced by high external focus in Experiment 1 of the present study (*N* = 104). Left panel illustrates bystander recall accuracy that follows a dosedependent function, with memory lowest for bystanders surrounded by 2 difficult discrimination trials, next lowest when adjacent to only 1 difficult discrimination, and best when the bystander was not near any difficult discriminations. Right panel illustrates growth of the amnesic shadow, expressed as the difference between 0 vs. 1 difficult epoch (orange) and 0 vs. 2 difficult epochs (dark red) conditions, changing over task quartiles. (B) For comparison, in prior work, retrieval stopping induced a comparable pattern of impaired memory formation and change over quartiles (data reproduced with permission from Hulbert et al.^65^, Experiment 1, *N* = 24). Error bars represent standard error of the mean. \**p <* .*05; ***p <* .*001*

Difficult discriminations demanded greater attentional resources than easy discriminations and may have distracted participants from processing upcoming bystanders. Such divided attention could reduce bystander encoding and impair memory. We tested this hypothesis with data from odd/even buffer trials presented between the discrimination and bystander tasks to account for effects of such distraction.

Firstly, a divided attention account predicts slower response times to buffer trials following difficult relative to easy discriminations, leading up to bystander presentation. However, reaction times were similar following difficult (*M* = 505ms, *SD* = 50.5) vs. easy (*M* = 508ms, *SD* = 57.1) discriminations (*F*(1, 101) = 0.43, *p* = .512, *BF*_*10*_ = .19). Buffer trial accuracy was higher following difficult (*M* = 92.8%, SD = 14.3) than easy (*M* = 90.4%, SD = 15.8) discriminations (*F*(1, 102) = 5.54, *p* = .021, *BF*_*10*_ = 2.15), contrary to the predictions of divided attention. Further analyses revealed that the buffer task was sensitive to detecting task-switching costs, which did not differ whether they followed difficult or easy discriminations when considering buffer data pooled across all three experiments reported here (see SI).

Secondly, divided attention predicts that story generation should be more challenging following difficult rather than easy discriminations. However, in the second half of the task (where the shadow dominated; see section entitled Growth over Presentation Quartiles), subjective difficulty ratings were not different for bystanders surrounded by zero (*M* = 2.04, *SD* = 0.65), one (*M* = 2.14, *SD* = 0.61), or two (*M* = 2.12, *SD* = 0.72) difficult discriminations (*F*(1.84, 176.61) = 1.86, *p* = .162, *BF*_*10*_ = .27). Likewise, time taken to make difficulty ratings was similar across zero (*M* = 2046ms, *SD* = 889), one (*M* = 2068ms, *SD* = 828), and two (*M* = 2114ms, *SD* = 902) epoch conditions (*F*(1.95, 187.58) = 1.43, *p* = .243, *BF*_*10*_ = .15). Therefore, we did not find evidence that participants were differentially distracted during the bystander task, as divided attention would predict. This result was observed across the three experiments reported here (see SI).

Together, buffer and bystander task performance was not reliably worse following difficult relative to easy discriminations, indicating no support for differential distraction from high external focus. These findings replicate results from the original hippocampal modulation paradigm using retrieval stopping^65,67,68^, where an amnesic shadow emerged without any accompanying evidence of distraction. Therefore, results from the buffer and bystander tasks do not support the possibility that divided attention induced bystander memory deficits.

### Anterograde and retrograde components

Previous research has found that bystanders located both before and after retrieval stopping were disrupted^65^, reflecting retrograde and anterograde effects. If difficult discriminations induce an amnesic shadow analogous to the one induced by retrieval stopping, both anterograde and retrograde components should arise. Firstly, including a single difficult discrimination either before or after a bystander impaired retention relative to including zero difficult discriminations (*F*(1, 96) = 5.44, *p* = .02, *η*_*p*_^*2*^ = .05, 90% CI = [.004, .14]). Moreover, bystanders flanked by difficult discriminations on both sides were recalled worse than bystanders flanked on only one side (*F*(1, 96) = 6.81, *p* = .011, *η*_*p*_^*2*^ = .07, 90% CI = [.009, .16]). Together these findings suggest a dose-dependent effect, because retention was lowest for bystanders surrounded by two epochs, next lowest for those near one epoch, and best for those near zero epochs (omnibus effect: *F*(1.92, 184.70) = 11.58, *p* < .001, *η*_*p*_^*2*^ = .108, 90% CI = [.04, .18]). Hence, including a difficult discrimination both before and after bystander presentation induced the greatest memory deficits, suggesting an additive contribution of both anterograde and retrograde effects.

Therefore, we observe that demanding external attention has a dose-dependent effect on bystander retention, mirroring amnesic effects observed by retrieval stopping^65^. The findings are consistent with additive contributions of both anterograde and retrograde components, suggesting that memory encoding and stabilisation processes may be disrupted by external focus. Moreover, differential distraction after difficult discriminations would not predict the involvement of retrograde components, providing further evidence that divided attention is not sufficient to explain the memory deficits.

### Growth over presentation quartiles

Previous research revealed that the amnesic shadow consistently grows over repeated retrieval stopping trials^65^, potentially due to greater below-baseline reduction of hippocampal activity over repeated suppression attempts^71^. If completing difficult discriminations induces an amnesic shadow analogous to the one induced by retrieval stopping, the shadow should similarly grow over quartiles in our attention modulation phase (see Figure 3). A linear growth of the shadow was found over presentation quartiles for bystanders surrounded by two, relative to zero, difficult discriminations (*β* = 1.88, 95% CI = [0.41, 3.36], *t*(96) = 2.53, *p* = .013). A similar growth was found for bystanders near one vs. zero surrounding difficult discriminations (*β* = 1.55, 95% CI = [0.28, 2.81], *t*(96) = 2.42, *p* = .017). Moreover, the shadow was observed in the second half of the attention modulation phase but not in the first half for the two vs. zero epoch comparison (interaction across halves: *F*(1, 96) = 16.4, *p* < .001, *η*_*p*_^*2*^ = .15, 90% CI = [.05, .26]; First half: *F*(1, 96) = 2.28, *p* = .134, *η*_*p*_^*2*^ = .02) and one vs. zero epoch comparisons (interaction across halves: *F*(1, 96) = 8.11, *p* = .005, *η*_*p*_^*2*^ = .08, 90% CI = [.014, .18]; First half: *F*(1, 96) = 2.95, *p* = .09, *η*_*p*_^*2*^ =.03).

Therefore, we find evidence that the amnesic shadow induced by external attention grows over quartiles, in line with previous observations of the amnesic shadow induced by retrieval stopping^65^.

### Selective impairment of source memory

Hulbert et al. observed that participants could recognise previously learned items despite being surrounded by retrieval stopping attempts, but the ability to recall the context in which those items were learnt was impaired^65^. These deficits in source memory suggested that retrieval stopping may specifically disrupt hippocampal-dependent episodic memory processes, aligning with dual-process models of recognition that implicate the hippocampus in forming traces containing episodic context rather than simply of the item^69,72,73^.

In our Experiment 2, we tested whether difficult discriminations would impair source memory but leave item recognition unaffected. The experimental paradigm (see Figure 4) and materials were similar to those used by Hulbert et al.^65^, except that difficult discriminations were used to induce an amnesic shadow instead of retrieval stopping. Participants were presented nouns, which were bystanders to surrounding easy and difficult discriminations, and asked to make either animacy (living vs. non-living) or pleasantness (pleasant vs. unpleasant) judgements. In a subsequent surprise recognition test, participants first rated their confidence of seeing a noun previously in the attention modulation phase (item recognition) and then recalled whether they had made an animacy or pleasantness judgement about that noun (source recognition). Based on prior evidence of the amnesic shadow and dual process models of recognition, we hypothesised that difficult discriminations would selectively impair source recognition, leaving item recognition intact, indicating a specific disruption to hippocampal-dependent episodic memory processing.

**Figure 4.**
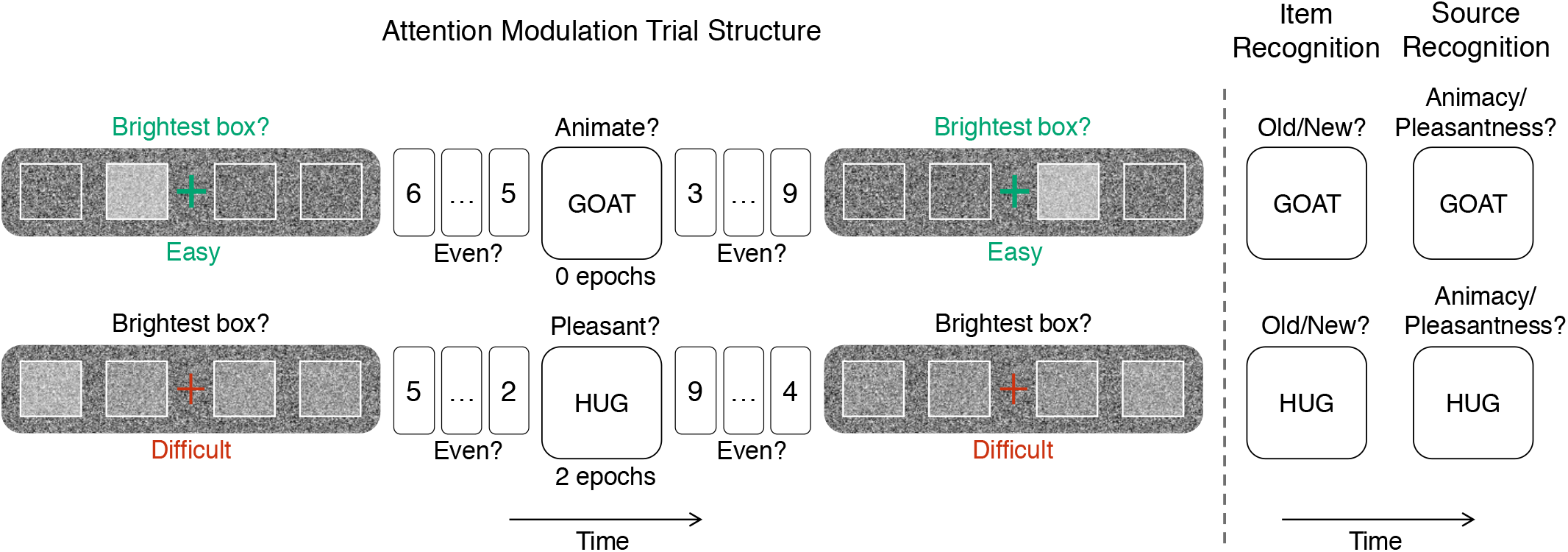
Trial structure of Experiment 2. In the attention modulation phase (left), nouns are presented as bystanders between easy (green; low external attention demand) or difficult (red; high external attention demand) discrimination trials, with odd/even buffer trials completed immediately before and after each bystander to account for task-switching effects. Participants make either an animacy (living vs. non-living) or pleasantness (pleasant vs. unpleasant) judgement for each noun. In the subsequent recognition test (right), participants first rate their confidence that they have seen the noun before (item recognition) on a scale from 1 (Definitely New) to 6 (Definitely Old) and then indicate whether they made an animacy or pleasantness judgement about that noun (source recognition).

Consistent with our predictions, we found deficits in source recognition for bystanders surrounded by two, relative to zero, difficult discriminations (*F*(1, 26) = 5.76, *p* = .024, *η*_*p*_^*2*^ = .18, 90% CI = [.015, .39], see Figure 5). However, item recognition did not reliably differ between zero (*M* = 80%, *SD* = 22) and two (*M* = 88%, *SD* = 21) epochs (*F*(1, 27) = 2.78, *p* = .11, *η*_*p*_^*2*^ = .09, 90% CI = [0, .29]), supported by a significant interaction between measures (*F*(1, 26) = 12.91, *p* = .001, *η*_*p*_^*2*^ = .33, 90% CI = [.10, .53]). We did not find a reliable dose-dependent effect (detailed report in SI). As in Experiment 1, the amnesic shadow for source recognition emerged in the second half of the experiment (reported above), but not in the first half (*F*(1, 28) = 0.88, *p* = .357, *η*_*p*_^*2*^ = .03; interaction with second half: *F*(1, 26) = 6.62, *p* = .016, *η*_*p*_^*2*^ = .20, 90% CI = [.02, .42]), suggesting that the shadow grows over time in the task. Item recognition did not reliably change across presentation halves (first vs. second half interaction: *F*(1, 27) = 0.88, *p* = .356, *η*_*p*_^*2*^ = .032). This selective emergence of the shadow in the second half, for source but not item recognition, was supported by a significant interaction (*F*(1, 26) = 6.97, *p* = .014, *η*_*p*_^*2*^ = .21, 90% CI = [.028, .42]). A numerical linear pattern was found across quartiles for source recognition (*β* = 0.53, 95% CI = [−0.15, 1.21], *t*(19) = 1.63, *p* = .12) and item recognition (*β* = −0.50, 95% CI = [−1.15, 0.14], *t*(19) = −1.65, *p* = .12), though neither was statistically reliable in isolation; the linear trend for source recognition significantly differed from that of item recognition (interaction: *t*(19) = 2.15, *p* = .044).

**Figure 5.**
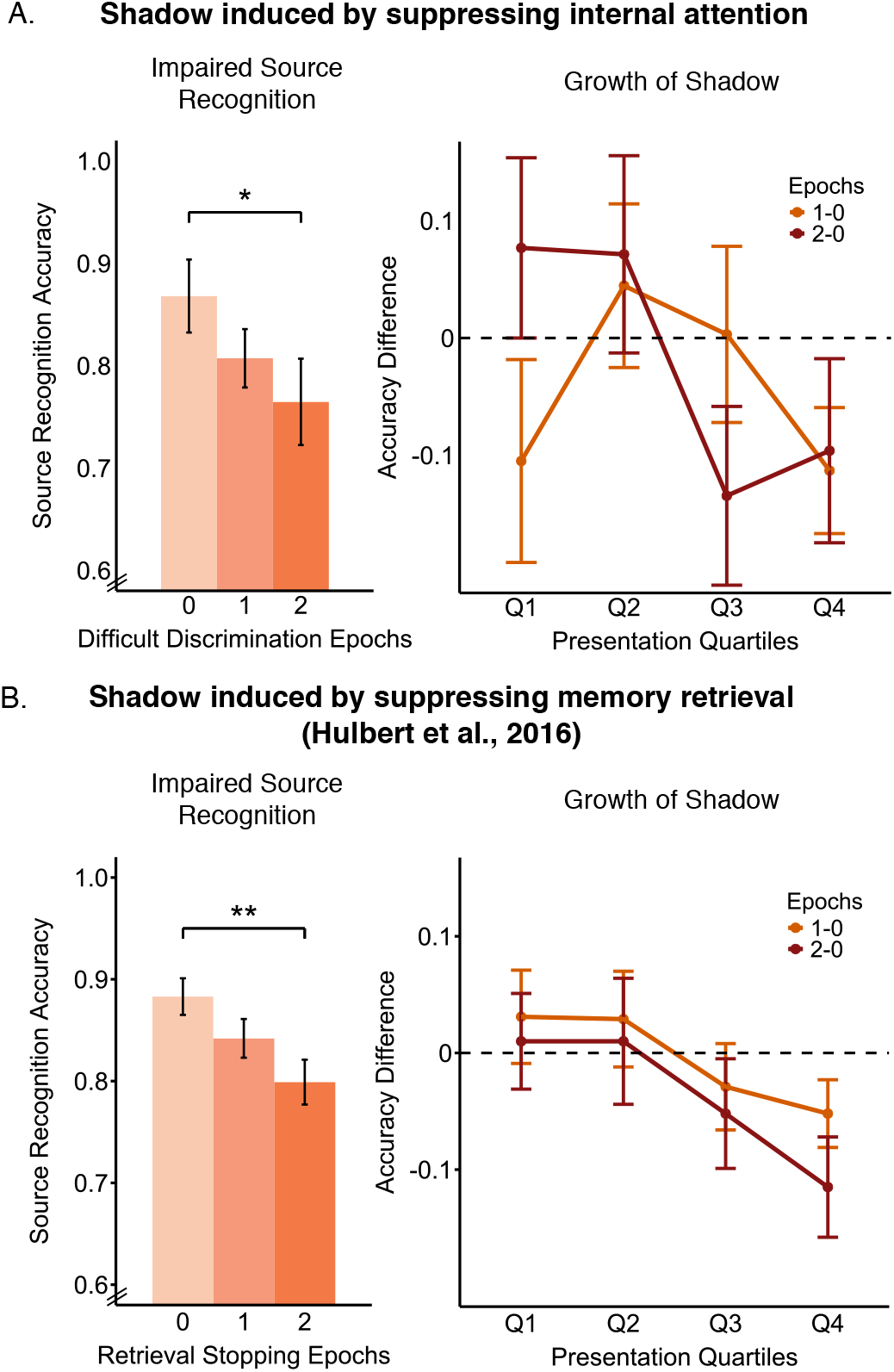
(A) The amnesic shadow for source recognition of bystander nouns, induced by demanding external focus in the present study (*N* = 30). Left panel illustrates source recognition accuracy, which was lower for bystanders surrounded by 2 difficult discrimination trials relative to bystanders surrounded by 0 difficult discrimination trials. Right panel illustrates numerical growth of the amnesic shadow over task quartiles, expressed as the difference between 0 vs. 1 epoch (orange) and 0 vs. 2 epochs (dark red) conditions. (B) Retrieval stopping induced a comparable pattern of deficits to source recognition and growth over quartiles (data reproduced with permission from Hulbert et al.^65^, *N* = 72). Error bars represent standard error of the mean. \**p <* .*05; **p <* .*01*

Overall, the results indicate that difficult discriminations selectively impair source recognition, which indexes the context in which a memory was formed, while no reliable deficits were found for the ability to recognise that item. This pattern is consistent with the possibility that hippocampal-dependent episodic memory formation is specifically impaired by difficult discriminations, similar to the patterns of the amnesic shadow observed during retrieval stopping^65^.

### Forgetting of previously formed episodic memories

Experiments 1 and 2 indicate that difficult discriminations impair the formation of novel episodic memories, demonstrating that episodic memory processing is reduced during high external focus. Two questions remain, however, regarding the mechanisms of these findings and their scope. First, as previously discussed, difficult discriminations could remove attentional resources from episodic memory processing and induce memory deficits. Although we do not find evidence in support of attentional withdrawal in prior experiments, the withdrawal vs. suppression hypotheses were not directly tested. Second, if difficult discriminations do cause amnesic effects, it remains unclear whether these effects are limited to the formation of new episodic memories or whether already encoded memories are also vulnerable to forgetting, in line with previous observations of the amnesic shadow^67,68^.

Experiment 3 addressed these questions by adapting a paradigm introduced by Zhu and Wang^68^. In the present paradigm (see Figure 6), participants learned associations between objects and negatively valenced scenes in a list, and between words and the same scenes in a separate list. The purpose of this learning phase was to first establish associations in long-term memory and then reactivate these memories between difficult discrimination trials, making already encoded memories bystanders to their potentially disruptive effects. Accordingly, during one bystander sequence in the subsequent attention modulation phase, a single object was presented between discrimination trials. The object was either previously learned with a scene (old) or shown for the first time (new). Participants indicated whether the object was old or new with a button press. Old objects were repeated across eight trial blocks, with new objects added in each block to maintain meaningful recognition judgements. As mentioned above, we assumed that presenting objects as reminders to previously encoded scenes would reactivate those scenes and render them susceptible to disruption^68^ from surrounding difficult discriminations, and that such disruption should not occur for scenes reactivated between easy discriminations.

**Figure 6.**
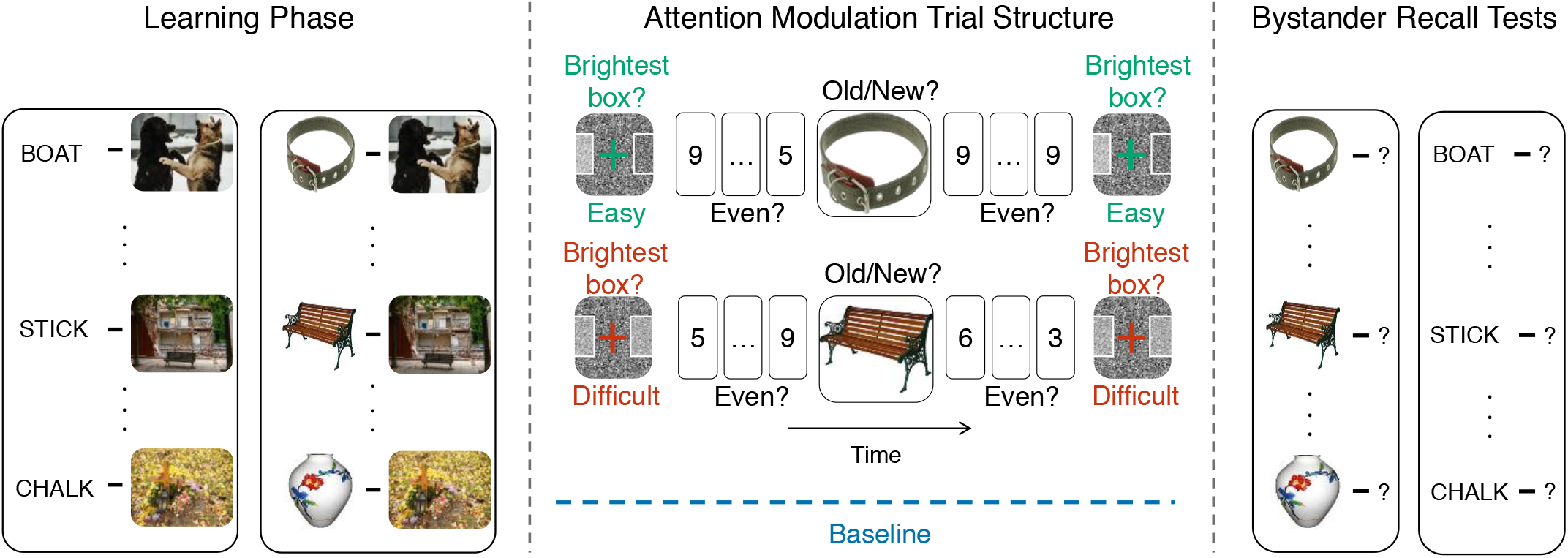
Trial structure of Experiment 3. In the learning phase (left), participants learned object-scene and word-scene associations as separate lists, with words and objects linked to the same scenes. In the attention modulation phase (middle), objects were presented between easy (green; low external attention demand) or difficult (red; high external attention demand) discrimination trials. The objects were either old and reactivated their associated scenes and made them bystanders to surrounding periods of external attention, or new. Participants indicated whether the object was old or new via button press. A baseline subset of scenes, previously learned with both objects and words, was never reactivated during the attention modulation phase (blue dashed line); baseline memories therefore had the same retention interval as the other memories but were unaffected by attentional modulation. In the bystander recall tests (right), scene memory was tested with both object and word cues; word cues were absent during the modulation phase and independent of any effects of completing the task, providing a direct measure of the scene memory traces. Example stimuli are for illustration purposes and were not used in the experiment.

Importantly, to test whether bystanders reactivated near difficult discriminations remained intact or were disrupted, we introduced a baseline subset. Baseline memories were learned at the same time as the reactivated memories but were not themselves reactivated. Retention of baseline memories therefore reflected passive forgetting over a matched interval, unaffected by easy or difficult discriminations. Participants thus learned three memory sets: one reactivated between difficult discriminations, one reactivated between easy discriminations, and a baseline set not reactivated at all. Comparing bystander retention against baseline distinguishes attentional withdrawal and active suppression: If difficult discriminations simply failed to provide a memory benefit, retention of their bystanders should match that of baseline memories suggesting that bystander memories are intact; if difficult discriminations actively disrupted bystanders, their retention should fall below that of baseline memories.

After the attention modulation phase, participants completed two surprise recall tests of scenes: one with object cues and another with word cues, which were both learned with the same scene prior to the attention modulation phase. The wordcue test addressed two key limitations of the object-cue test, one theoretical and one methodological. Theoretically, forgetting of scenes measured in the object test could reflect disruptions to the retrieval route used by objects to reactivate the bystander scenes, or to the scene itself. Probing scene memory with objects would not distinguish these possibilities. Since the word cues were never shown in the attention modulation phase, any specific disruptions to the scene memory should be detectable by these independent cues^70,74^. Hence, if forgetting generalises to the word cue test, then it would indicate disruptions to the bystander memory itself, not just the retrieval route used by object cues. Methodologically, the word cues were also independent of an extraneous source of forgetting introduced by using object cues to reactivate the scenes. Repeatedly viewing objects while making old/new judgements could create new associations between the object and the task rule (responding “old”) which could interfere with the retrieval route of the object cue to the scene. Such associative interference^70^ could impair bystander memory below baseline for both easy and difficult discriminations when probing memory with the object cues, adding to and obscuring any specific disruptive effects of difficult discriminations. The word cues thus provided a measure of specific disruptions to scene memories themselves, independent of associative interference.

Scene memory from the bystander recall tests was quantified using two measures: identification, which assessed whether participants provided unambiguous descriptions of the scenes; and gist, which measured recall of a predefined set of central scene details^109^. The results presented below followed a preregistered analysis plan (see OSF for details). Results are illustrated in Figure 7.

**Figure 7.**
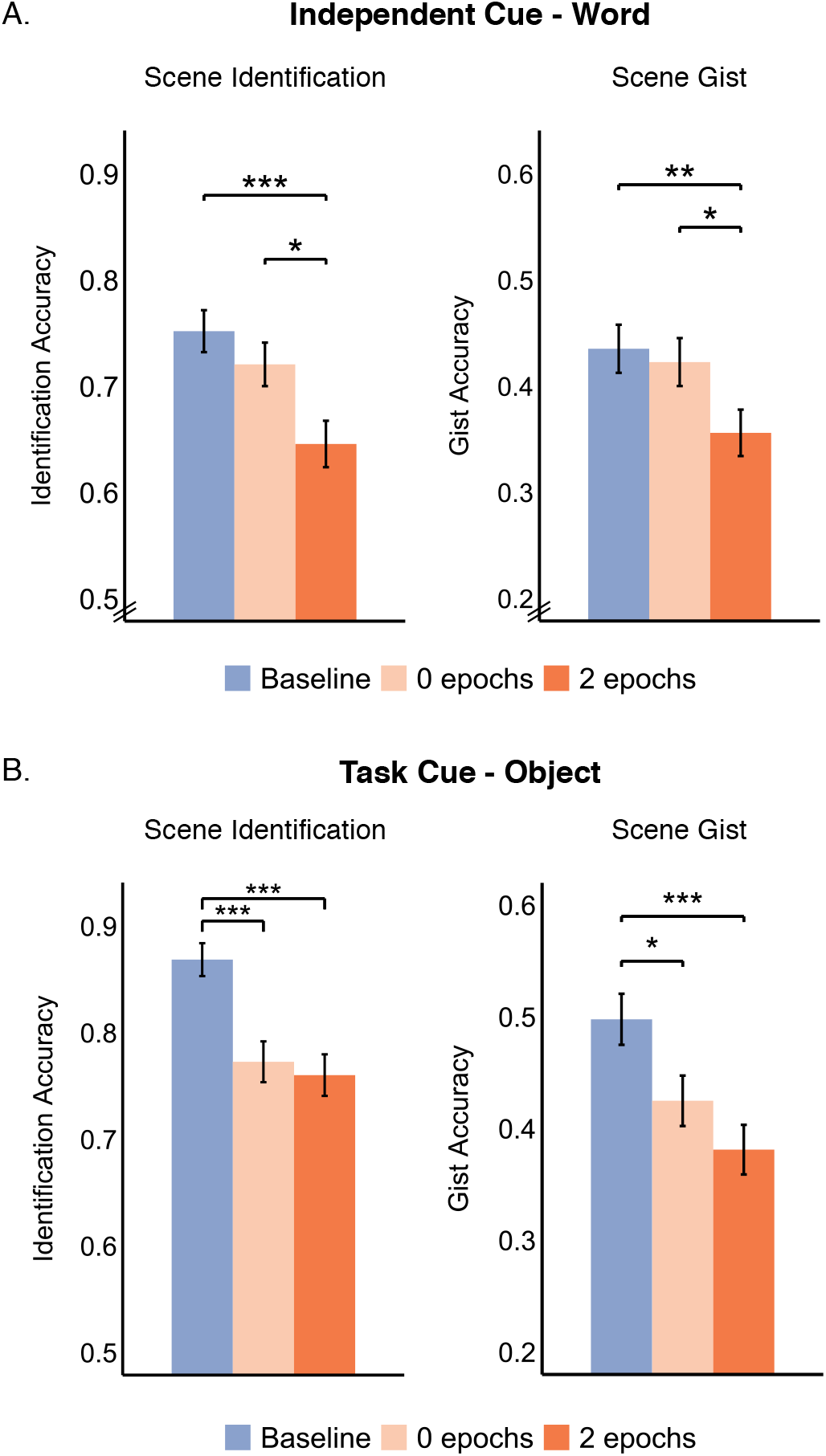
The amnesic shadow for already encoded episodic memories reactivated near demanding external focus in Experiment 3 (*N* = 60). Left panels show scene identification accuracy and right panels show scene gist accuracy, each as a function of attentional demand: baseline memories that were not subject to any attentional demand (blue), bystanders reactivated near zero difficult discrimination epochs (light orange), and bystanders reactivated near two difficult discrimination epochs (dark orange). Scene memory was tested with both task-independent word cues (A) and object cues used to reactivate scenes in the attention modulation phase (B). (A) Word-cued recall was selectively worse for bystanders reactivated near two difficult epochs relative to both baseline and zero epochs, while zero epochs did not reliably differ from baseline, demonstrating disruption to scene memory traces. (B) Object-cued recall was worse for both zero and two difficult epochs relative to baseline, possibly due to a combination of the amnesic shadow for difficult discriminations, and extraneous associative interference from task instructions unrelated to attentional demand. Error bars represent standard error of the mean. *\*p <* .*05, **p <* .*01, ***p <* .*001*.

We first aimed to extend the findings from Experiments 1 and 2, which indicated that nearby difficult discriminations impair the formation of episodic memories relative to easy discriminations. If already encoded memories are similarly vulnerable, scenes reactivated between difficult discriminations should be recalled worse than those reactivated between easy discriminations. The evidence supported this prediction across cue types (identification: *F*(1, 57) = 4.81, *p* = .032, *η*_*p*_^*2*^ = .08, 90% CI = [.004, .21]; gist: *F*(1, 57) = 5.54, *p* = .022, *η*_*p*_^*2*^ = .09, 90% CI = [.007, .22]). This forgetting effect did not reliably interact with cue type (identification: *F*(1, 57) = 3.42, *p* = .07, *η*_*p*_^*2*^ = .06; gist: *F*(1, 57) = .44, *p* = .51, *η*_*p*_^*2*^ = .008), supporting generalisation across object and word cues. Therefore, the evidence indicates that the amnesic shadow observed in Experiments 1 and 2 extends to already encoded memories.

The foregoing findings do not clarify whether difficult discriminations simply deprived bystanders of the benefits of attention to memory relative to easy discriminations, as predicted by attentional withdrawal. In such a case, bystanders should remain intact and their retention should not fall below that of baseline memories. However, if retention falls below that of baseline memories, it would indicate active disruption of bystanders, as predicted by active suppression. Supporting active suppression, we found below-baseline forgetting of bystanders near difficult discriminations (identification: *F*(1, 57) = 36.98, *p* < .001, *η*_*p*_^*2*^ = .39, 90% CI = [.23, .53]; gist: *F*(1, 57) = 21.89, *p* < .001, *η*_*p*_^*2*^ = .28, 90% CI = [.13, .42]). Hence, difficult discriminations did not merely limit the enhancement of bystander memories — they disrupted their retention. Moreover, the below-baseline forgetting of bystanders surrounded by difficult discriminations did not interact with cue type (identification: *F*(1, 57) = .005, *p* = .95, *η*_*p*_^*2*^ < .001; gist: *F*(1, 57) = 1.61, *p* = .21, *η*_*p*_^*2*^ = .03), hence generalising to the independent cue, and indicating disruptions to the scene memories themselves.

A remaining concern was that associative interference from old/new judgements on the objects could induce below-baseline forgetting of bystanders to both difficult and easy discriminations (see SI for results supporting this concern). Word cues were unaffected by such task-based influences, providing a test that isolates the selective disruptive effect of difficult discriminations. Specifically, forgetting of bystanders near difficult discriminations should exceed that of both baseline memories and bystanders near easy discriminations. However, we should not observe below-baseline forgetting for bystanders near easy discriminations. Such patterns would indicate that difficult discriminations selectively disrupt the scene memories themselves by ruling out extraneous interference effects.

Results from the word cue test suggested selective disruption to scene memories by surrounding difficult discriminations. Bystanders near difficult discriminations were recalled worse than baseline scenes (identification: *t*(59) = 4.13, *p* < .001, *d* = .42, 95% CI = [.21, .64]; gist: *t*(59) = 2.93, *p* = .005, *d* = .33, 95% CI = [.10, .57]), and worse than bystanders near easy discriminations (identification: *t*(59) = 2.53, *p* = .014, *d* = .31, 95% CI = [.06, .56]; gist: *t*(59) = 2.21, *p* = .031, *d* = .28, 95% CI = [.02, .54]). Moreover, recall of bystanders between easy discriminations did not reliably differ from baseline (identification: *t*(59) = 1.13, *p* = .26, *d* = .13; gist: *t*(59) = .40, *p* = .69, *d* = .05). These patterns were supported by a significant omnibus effect across the three discrimination difficulty conditions (baseline vs. easy vs. difficult for identification: *F*(1.93, 110.24) = 7.66, *p* < .001, *η*_*p*_^*2*^ = .12, 90% CI = [.03, .21] and gist: *F*(1.94, 110.67) = 4.22, *p* = .018, *η*_*p*_^*2*^ = .069, 90% CI = [.007, .15]).

Together, these results indicate that demanding external focus disrupted already-encoded scene memories that were reactivated in their temporal vicinity.

## Discussion

Our findings suggest that focusing on the external world actively suppresses our thoughts, rather than merely depriving them of attention. Thoughts about the past, future plans, worries, and stray ideas that are unrelated to the task may interrupt external focus^20,26^, and may need to be suppressed. Such thoughts may spontaneously arise from replay mechanisms^55,59^, which depend on internally directed processing centred on the episodic memory system^40,52,53,75^. Hence, external focus may include the suppression of episodic memory processes.

Three experiments tested whether periods of high external focus disrupted episodic content, a measurable index related to internally directed thoughts. In Experiment 1, difficult visual discriminations impaired memory formation of novel episodic content presented near in time to the discriminations, relative to easy discriminations. Experiment 2 revealed specific deficits to the episodic context in which memories were formed, rather than retention of the item itself. In Experiment 3, previously encoded memories reactivated near difficult discriminations were forgotten below baseline memories unaffected by any attentional modulation and remained inaccessible even when cued with an independent cue. Together, demanding external focus disrupted memory for unrelated ‘bystander’ content near in time, casting an ‘amnesic shadow’. Hence, the amnesic shadow indicates that our thought contents are disrupted as a consequence of turning attention outward.

Our evidence for an amnesic shadow is consistent with an active suppression mechanism rather than divided attention or attentional withdrawal. In principle, difficult visual discriminations could fatigue participants, dividing attention away from processing imminent bystander content and reducing its retention. This divided attention hypothesis predicts worse performance on the odd/even buffer and bystander tasks following difficult relative to easy discriminations. Our data from buffer and bystander tasks did not follow this pattern. The attentional withdrawal hypothesis proposes that difficult discriminations require greater removal of attention from memory than easy discriminations, pausing memory processes that would otherwise benefit bystander retention. Indeed, difficult discriminations reduced bystander retention relative to easy discriminations across experiments. However, in Experiment 3, bystander retention was reduced below that of baseline memories that were not reactivated between visual discriminations. Merely depriving attention from bystanders cannot explain such below-baseline forgetting. Likewise, simply reducing bystander processing should also not produce below-baseline forgetting, further questioning the sufficiency of the divided attention hypothesis. Therefore, neither divided attention nor attentional withdrawal explains the observed amnesic shadow.

Two behavioural signatures of active suppression were observed here. First, active suppression may be involved if forgetting exceeds what is expected from the passage of time alone^63,64,68^. In Experiment 3, passive forgetting over time was measured with the retention of baseline memories. Therefore, the observed below-baseline forgetting in Experiment 3 likely reflects an active process. Second, active suppression could degrade the memory itself, beyond disrupting a specific retrieval route used to access the memory, exhibiting cue independence^70^. Crucially, we found below-baseline forgetting even when probed with an independent cue, indicating that the memory itself was degraded. The observed amnesic shadow therefore exhibits both below-baseline forgetting and cue-independence, which together point to the involvement of active suppression^63,64,67,68,70^. External focus therefore appears to recruit an active suppression mechanism that degrades thought contents, rather than merely depriving them of attention via attentional withdrawal.

How does external focus suppress thought contents? We consider two possibilities. External focus could broadly suppress the processes that give rise to distracting thoughts, stopping such content from arising at its root. Alternatively, individual thoughts that surface could be selectively targeted. One situation where selective suppression could occur is when thought contents share semantic features with perceptual stimuli, directly competing for prioritisation. Mutual inhibition would selectively suppress such content to prioritise perception^76,77^. Supporting the existence of such a mechanism, a previous study found that perception and memory regions show opposing responses at matched retinotopic locations when perceiving scenes relative to recalling scenes from memory^78^, displaying mutual inhibitory dynamics. However, our bystander stimuli comprised diverse content largely unrelated to the low level perceptual stimuli of the visual discrimination task. These findings suggest that episodic memory processing is broadly suppressed, disrupting any episodic content that is active near in time. Such systemic suppression is consistent with previous observations of the amnesic shadow induced by retrieval stopping, where diverse unrelated episodic content was forgotten^64,65,67,68^. The present findings further support the possibility of systemic suppression of episodic memory processing, a newly discovered source of forgetting. Together, the evidence suggests that external focus may inhibit episodic memory processes globally, rather than selectively targeting thought contents.

One plausible means to inhibit episodic memory processing is to suppress hippocampal activity. The hippocampus is known to support the formation of novel memories, retention of episodic context, and the replay and consolidation of already encoded memories^79–81^. Suppressing hippocampal activity may therefore be a basis for globally inhibiting these processes. Supporting this possibility, intentionally stopping memory retrieval suppresses hippocampal activity^40^, substantiated by reduced hippocampal BOLD activity^62,82^ and theta oscillations^83–85^, with higher hippocampal GABA concentrations predicting greater forgetting of those memories^86^. Such retrieval stopping induces an amnesic shadow for unrelated bystanders^64,65,67,68^, with the magnitude of this shadow scaling with hippocampal BOLD reductions^65^. Hippocampal suppression may therefore underlie the systemic disruption of episodic memory during retrieval stopping. Crucially, demanding sensory attention reliably decreases hippocampal activity^3,61^. Specifically, a previous study used the same visual discrimination task as was used here and observed lower hippocampal BOLD activity during difficult relative to easy visual discriminations^61^. It is possible that lower hippocampal activity during external focus reflects hippocampal suppression. Therefore, although neural activity was not measured here, we speculate that the observed amnesic shadow is a behavioural manifestation of hippocampal suppression during external attention.

The target of suppression may extend beyond the hippocampus to the default network, which supports internally directed cognition more broadly. The hippocampus communicates with the default network to support episodic memory^2,3,87,88^, particularly within DN-A, a subsystem of the default network specialised for episodic memory processing^87,89–91^. External sensory attention reliably reduces default network activity^11–17^. Hence, the default network may also be a target of systemic suppression during external attention. Reductions in default network and hippocampal activity have remained functionally ambiguous^3,19^, because such reductions could reflect passive disengagement caused by truncated input or active suppression via an inhibitory signal. The observed amnesic shadow is consistent with the possibility that the default network is often suppressed, rather than passively disengaged, during sensory attention. Nevertheless, systemic suppression of hippocampal and default network activity would benefit external focus by stopping replay mechanisms, which drive spontaneous thoughts and mindwandering^7,54,55,58–60^.

Suppressing the cognitive processes that support replay may therefore protect external attention by stopping thought contents from becoming active and triggering attentional capture back inward. This is a novel mechanistic proposal for stopping internal distraction during external attention, distinct from previous accounts. Attentional capture could instead be prevented by suppressing the orienting mechanism that pulls attention inward, so that internal contents fail to interrupt external focus even when active. Precedent for this idea comes from proposals of how external distraction is handled: salient external stimuli generate a priority signal that captures attention away from the goal-relevant stimulus, and this signal can be actively suppressed to protect task engagement^37,92–94^. Because episodic memories may compete for attentional resources with external stimuli^8,32,43–49,95,96^ by engaging similar capture mechanisms^40,42^, suppressing attentional orienting could stop capture by internal content as it does for external content. Consistent with this possibility, activity in the ventral attention network, on which attentional capture depends^35^, is reduced during external focus^97–99^. However, only suppressing attentional orienting should not affect memory, leaving memory traces intact rather than degraded as observed here. If attentional orienting is suppressed during external focus, it must therefore operate alongside, rather than instead of, the suppression of episodic memory systems.

Focusing on the external world may spontaneously suppress our thoughts. Here, participants intended to discriminate brightness levels of visual stimuli, without an explicit goal to suppress memory. Yet, bystander memories active close in time to difficult discriminations were disrupted. Systemic suppression of memory processing has been observed only during the deliberate stopping of memory retrieval^40,63,64,100^. Our findings therefore provide novel evidence for the incidental engagement of process suppression. Interestingly, the observed amnesic shadow shares several properties with the shadow induced by retrieval stopping^65,67,68^: a dose-dependent effect, combination of anterograde and retrograde components, growth over repeated trials, impaired memory formation, specificity to episodic context, below-baseline forgetting, and cue independence. Hence, we speculate that external focus may spontaneously engage a suppression mechanism similar to the one that is intentionally engaged during retrieval stopping. Retrieval stopping involves a top-down ‘stop’ signal from the right anterior dLPFC that suppresses the hippocampus^40,101,102^, and a similar signal could therefore be implemented during external focus. Alternatively, the right anterior insula may be involved, as it detects salient stimuli and initiates suppression of the default network^19,103–106^. However, the neurocognitive mechanisms underlying the cascade from initiation via the insula to termination of default network activity remain unspecified. Regardless of the precise mechanism, our findings suggest that attending to the external world may spontaneously suppress the cognitive processes that generate distracting thoughts.

In conclusion, our findings challenge the idea that focusing on the external world simply withdraws attention from our thoughts. Rather, external focus can spontaneously recruit an active suppression mechanism that may act broadly on processes supporting the emergence of spontaneous thoughts, stopping such thoughts from capturing attention and interrupting focus. As a consequence, our thoughts may not be merely paused when attention turns outward, but actively degraded. By suppressing internal distraction in this way, this mechanism may keep attention anchored to the external world. Notably, other foci of internal attention such as pain and other interoceptive signals^107,108^ may also be sources of internal distraction that could be suppressed during external attention. Our findings also speak to clinical conditions marked by persistent thoughts, where external attention is often poorly sustained^28–31^. If external focus ordinarily suppresses such thoughts, difficulty in sustaining external focus may itself reflect a failure of the suppression mechanism. More broadly, the findings provide a novel proposal for how internal distraction can be regulated in daily life: We may unknowingly suppress our thoughts by simply turning attention to the external world.

## Methods

### Experiment 1

#### Participants

One hundred and four adults between ages 18 and 52 years old (M_age_ = 24.47 years, SD = 7.76, 55 females and 49 males) participated. The sample included pooled data from two datasets (termed 1a and 1b), collected across the University of Oregon, USA; University of St. Andrews, Scotland, UK; and MRC Cognition and Brain Sciences Unit, University of Cambridge, UK. All participants had normal or corrected-to-normal vision, had no prior or existing neurological or psychological illnesses including learning, reading, or attention disorders. From dataset 1a, six participants were excluded as they reported sleeping less than 6 hours during the night before participating in the study, and one of these participants also had ADHD. Twelve further participants from this dataset were excluded because they could not complete the staircasing procedure. There were no participant exclusions in dataset 1b. Participants were recruited through university course-credit schemes, word of mouth, or public fliers, and received course credit or monetary compensation.

Dataset 1a included 56 participants, collected before widespread adoption of preregistration practices, exceeding typical sample sizes of the time (*N* ~ 30). Dataset 1b aimed to replicate the primary amnesic shadow effect (zero versus two-epoch difference), and included an a priori sample size of 48 (see OSF link for preregistration). To maximise statistical power, both datasets were pooled. A sensitivity analysis on the combined sample (*N* = 104) indicated 80% power to detect an effect size of *d* = 0.27 for a paired t-test (two-tailed, *α* = .05).

#### Task Description

The attention modulation phase comprised three tasks, described below. The procedure for the attention modulation phase is described in the Procedure and Design section.

##### Visual discrimination task

A visual discrimination task was used to modulate demand on external attention (see Figure 2). The task parameters closely matched a previous study that observed lower hippocampal activity in the difficult relative to easy version of this task^61^.

In one visual discrimination trial, a fixation cross was presented on a black background for 750ms, followed by four boxes presented horizontally against static grayscale noise for 1200ms. One box (target) was brighter than the other three (lures). Then, a “Go!” cue replaced the fixation cross, prompting participants to indicate the brightest box via button press within 700ms. In addition to highlighting the correct response option, “Yes” was presented as feedback if participants selected the target box; “No” was presented if they selected a lure or did not respond within 700ms. Feedback appeared as soon as a response was received and remained on screen for the remainder of the 700ms response window. The feedback was then presented for an additional 800ms. A fixation cross then reappeared for 250ms to close out the trial.

In the easy version, the target was 50% brighter than the three lures, which were fully transparent. In the difficult version, the target was marginally brighter than the lures (*M* = 8.17%; *SD* = 4.61). For each difficult trial, lures were randomly assigned a single brightness value between 0 (fully transparent) and 50% (half-opaque). The fixation crosses, “Go!” cue, and feedback were coloured green for easy trials and red for difficult trials, allowing participants to prepare for the upcoming difficulty.

We calibrated the difficulty of difficult discrimination trials for each participant such that they would correctly identify the target box on approximately 50% of trials, in line with Law et al.^61^. An adaptive staircasing task was conducted before the attention modulation phase. The exact staircase procedure differed somewhat between datasets 1a and 1b, but overall discrimination performance did not significantly differ across datasets.

##### Bystander Task

Bystanders were images of an object embedded in a scene. We used the same set of bystander images as Hulbert et al.^65^. In one trial, a pre-cue “Object/Scene” was presented for 500ms, followed by the image for up to 5000ms. Participants answered the question “Why is the pictured object in this location?” and rated the difficulty in generating a story on a scale of 1 (Not difficult at all) to 4 (Extremely difficult). Hence, the task encouraged associative encoding of the object and scene. The trial ended as soon as a response was recorded.

##### Buffer Task

Buffer tasks were included to control for task-set switching costs between discrimination and bystander tasks. An “even?” warning was presented in white on a black background for 500ms, followed by a digit (1–9) for 900ms, during which participants indicated whether the number was even (Yes) or odd (No). Between 2 and 4 digits were presented per task (except for non-filler trials at the beginning and end of each block, which had more relaxed constraints), with a 400ms inter-stimulus interval between each digit. Each trial ended as soon as a response was recorded. The specific digits presented and the number of digits per task were predetermined and identical across participants.

#### Procedure and Design

Participants initially practised each task individually, and then combined together as part of a single sequence, as they would appear in the attention modulation phase. Then, they completed the crucial attention modulation phase, in which bystanders were presented within a stream of visual discriminations. Finally, they completed a surprise bystander memory test.

##### Attention Modulation Phase

Participants completed a stream of easy and difficult visual discrimination trials, occasionally interrupted by bystander images. When a bystander was presented, the task sequence followed the pattern: visual discrimination task, buffer task, bystander task, buffer task, visual discrimination task (see Figure 2), referred to as a bystander sequence. A 400ms ITI separated each task within the bystander sequence.

Thirty-two bystander images were divided equally into four sets, and each set was assigned to a condition determining the bystander’s placement before or after a difficult discrimination trial. The four conditions were: Bystander presented i) After difficult and before difficult; ii) After difficult and before easy; iii) After easy and before difficult; iv) After easy and before easy. Set assignment was counterbalanced across participants in both datasets. In dataset 1b, assignment of bystanders into different quartiles of the attention modulation phase was additionally counterbalanced across participants.

The presentation order was pseudorandomised prior to data collection and remained constant across all participants.

##### Surprise Bystander Memory Test

After the attention modulation phase, participants were tested on their memory for bystanders. In each trial, the background scene from one bystander appeared for 15s without the embedded object. Participants recalled and verbally reported the object they had seen in that scene during the attention modulation phase. The experimenter rated response accuracy based on preregistered rules for 1b (see OSF).

### Experiment 2

#### Participants

Thirty adult participants (ages 18–26 years; *M*_*age*_ = 20.37 years, *SD* = 1.74; 21 females, 9 males) were included in this experiment. Data were collected at the University of Oregon, USA and University of St. Andrews, Scotland, UK. There were no participant exclusions, although some analyses excluded participants with no remaining trials in a condition (see Statistical analysis approach). All participants were neurologically healthy without diagnosed learning, reading, or attention disorders, and had normal or corrected-to-normal vision. All participants were exposed to English as a primary language since early childhood (0–3 years). Participants were recruited through university course-credit schemes, word of mouth, or public fliers, and received course credit or monetary compensation.

The sample was collected in 2009, prior to widespread adoption of preregistration practices, and matched typical sample sizes of that time. A sensitivity analysis indicated 80% power to detect an effect size of *d* = 0.53 for a paired ttest (two-tailed, *α* = .05).

#### Task Description

##### Visual discrimination task

Visual discrimination task parameters closely matched the task used in Experiment 1, with one exception that the precue fixation cross was presented for 300ms instead of 750ms.

##### Bystander Task

Bystanders were the same nouns as used in Hulbert et al.’s Experiment 6a^65^. In one trial, a precue (“pleasant?” or “living?”) was presented for 500ms, followed by the noun for 3500ms. Participants judged the animacy (living or non-living?) of half the bystanders and the pleasantness (pleasant or unpleasant?) of the other half, counterbalanced across participants. The trial ended as soon as a response was received.

##### Buffer task

The buffer task followed the same trial structure as Experiment 1, but participants completed as many odd/even judgements as possible within a fixed 5000ms window. Digits were randomly generated by the presentation software on each trial and were separated by 100ms intervals.

#### Procedure and Design

##### Attention Modulation Phase

The attention modulation phase followed the same procedure as Experiment 1, and included a similar task structure: visual discrimination task, buffer task, bystander task, buffer task, visual discrimination task. A 500ms interval separated each task within the bystander sequence. Participants completed the attention modulation phase with visual discriminations, as well as the Think/No Think version of the task across two separate blocks, in one session. The block order was counterbalanced across participants (TNT first or discrimination first). Only data from the attention modulation phase are reported here due to its relevance to the research question.

A total of 64 nouns were used and counterbalanced across 8 groups between the Think/No-Think and visual discrimination versions. For the purposes of this article, thirty-two bystander nouns presented in the discrimination version were assigned into the four conditions as in Experiment 1: Bystanders presented i) After difficult and before difficult; ii) After difficult and before easy; iii) After easy and before difficult; iv) After easy and before easy.

The presentation order was pseudorandomised prior to data collection and remained constant across all participants.

##### Surprise Bystander Recognition Test

Participants completed a surprise recognition test for bystander nouns. The test included the 32 bystander nouns (targets) alongside 32 nouns not previously seen in the experiment (foils). In each trial, a noun appeared for 15s and participants reported their confidence in having seen that word previously on a scale of 1 (Definitely New) to 6 (Definitely Old), measuring item recognition. In that same trial, participants indicated whether they had completed the animacy or pleasantness judgement on that noun during the attention modulation phase within 15s, measuring source recognition.

### Experiment 3

#### Participants

Sixty human adult participants aged between 18 and 43 years (*M* = 26.45 years, *SD* = 5.91, 43 females and 17 males) were included in this experiment. Data were collected at the MRC Cognition and Brain Sciences Unit, University of Cambridge, UK. All participants had normal or corrected-to-normal vision, no neurological or psychological illnesses, and were not on any psychoactive drugs. The experiment was terminated for six participants who could not learn 100% of the scene associations during the initial learning phase. Two additional participants were excluded as outliers: one based on visual discrimination task performance and one based on odd/even task performance. A priori power analyses indicated 83% power to detect an effect of *d* = .38 for a paired t-test (two tailed, α = .05). The study design, hypotheses, sample size, and exclusion criteria were preregistered. Participants were recruited through the MRC CBU’s SONA system, word of mouth, or public fliers, and received monetary compensation.

#### Task Description

The visual discrimination and buffer tasks were identical to Experiment 1.

##### Bystander Task

Bystanders were negatively valenced scenes, each paired with an object and a word across separate learning lists. The objects and scenes were selected from stimuli curated by Küpper et al.^109^. The words were selected from the MRC Psycholinguistic Database^110^; only nouns rated Medium (400) to High (700) on Familiarity, Concreteness, Imageability, and Meaningfulness were included.

In one trial, a precue “Old?” was presented for 500ms, followed by an object for 2000ms. Participants indicated whether they had seen the object before (“Old”) or not (“New”) via a button press within 2000ms. We assumed that objects previously learned with scenes would reactivate the scene, making the scene a bystander to surrounding discrimination tasks. The trial ended immediately after a response was recorded. Word cues were never presented during the attention modulation phase, providing a task-independent retrieval cue at test.

#### Procedure and Design

##### Learning and Test-Feedback Phase

Participants learnt each bystander scene with an object in one list, and a word in a separate list. In each trial, an object was presented on the left and a scene on the right with a fixation cross in the centre of the screen for 6000ms, and participants had to silently associate the scene with the object. There was a 1000ms inter-trial interval between trials.

After learning this list, participants were tested on their knowledge of the object-scene pairs. Each trial started with a fixation cross shown for 750ms. Then, an object was presented by itself for 4000ms, and participants indicated whether they could recall the associated scene via a button press (Yes or No) during that time. If “Yes,” they completed a 3-AFC recognition test for the scenes within 4000ms with the object appearing on top of the screen. If they picked the wrong scene, “WRONG” was presented for 300ms. Then, the original object-scene pair was presented as feedback for 2500ms, regardless of retrieval success. Participants who did not learn 100% of the object-scene pairs within three attempts did not continue beyond this stage of the experiment, as preregistered (see OSF). The same process was completed for a list of wordscene associations. The experiment was also terminated if participants did not learn 100% of word-scene cues. Participants were encouraged to create unique object-scene and word-scene associations, which aimed to avoid the possibility of integrating the object, scene, and word into one coherent memory^111–113^.

##### Attention Modulation Phase

Unlike Experiments 1 and 2, every trial in the attention modulation phase included a bystander. Each trial followed the same bystander sequence structure: visual discrimination task, buffer task, bystander task, buffer task, visual discrimination task. A 2000ms ITI separated each bystander sequence. The presentation order was pseudorandomised prior to data collection and remained constant across all participants.

The attention modulation phase comprised 8 blocks. Each block included 16 previously learned objects (“Old”; 8 easy, 8 difficult) and 8 foils (“New”) presented as bystanders. A new set of 8 foil objects was used in each subsequent block, giving 64 foils and 16 old objects across the 8 blocks.

A total of 24 scene-object-word triads were divided into three sets and assigned to a condition based on their placement between difficult discriminations: 1) Zero epochs, where the bystander was flanked by two easy discriminations; 2) Two epochs, where the bystander was flanked by two difficult discriminations; and 3) Baseline, scenes previously learned but not reactivated during the attention modulation phase. Assignment of stimuli into the three conditions was counterbalanced across participants.

##### Surprise Bystander Memory Test

Participants’ scene memory was tested first using objects in one test list, and then with words in a separate test list (or vice versa, as list order was counterbalanced across participants). On each trial, the object or word cue was displayed for 15s, during which participants verbally described the associated scene in as much detail as possible. Scene descriptions were transcribed and scored on gist and identification measures described in the statistical analysis approach section. Trials were separated by a 3s ITI.

#### Statistical analysis approach

All experiments followed a repeated measures design. The primary independent variable was the number of difficult discrimination epochs surrounding the bystander. The dependent variable was bystander recall accuracy measured during the surprise memory test.

Our analysis largely followed the approach by Hulbert et al.^65^. In the 0 epoch condition, bystanders were flanked by two easy discriminations. In the 1 epoch condition, bystanders were flanked by one easy and one difficult discrimination. In the 2 epoch condition, bystanders were flanked by two difficult discriminations. Experiment 3 did not include a 1-epoch condition; bystanders were assigned to 0 epochs, 2 epochs, or a Baseline condition (learned but not presented during the attention modulation phase).

##### Experiments 1 and 2

In Experiment 1, cued-recall accuracy was analysed in a mixed ANOVA with epoch condition (0, 1, 2) and task half (first, second) as repeated measures factors. A parallel analysis was conducted for anterograde (difficult discrimination before bystander) and retrograde (difficult discrimination after bystander) effects, using the 1-epoch condition split by direction. Dataset (1a vs. 1b) was included as a nuisance factor.

In Experiment 2, responses along the six-point confidence scale (1 = Definitely New; 6 = Definitely Old) were binarised into “new” (1–3) or “old” (4– 6). Item recognition accuracy was computed by comparing reported status (old or new) to veridical status (target or foil). Source recognition accuracy was scored as correct (1) or incorrect (0) based on whether participants accurately identified the judgement type (animacy or pleasantness) completed for that noun. Only correctly recognised items were further an alysed for source recognition. Both measures were analysed in a mixed-design ANOVA with epoch condition (0, 1, 2), recognition measure (item, source), and task half (first, second) as repeated measures factors. The task order (TNT or Discrimination version completed first) was included as a nuisance factor, because the retention interval from discrimination task to bystander recognition test significantly varied between the two counterbalancing groups.

The amnesic shadow was defined as the difference in cued-recall (Experiment 1) or source recognition (Experiment 2) accuracy between the zero and two-epoch conditions, for bystanders presented in the second half of the attention modulation phase^65^.

Missing responses were excluded before computing means from the remaining trials. Participants with no remaining trials in a condition were removed from the affected analysis (listwise deletion). Trials were missing mainly in dataset 1a and Experiment 2, owing to technical errors during data collection. In addition, source recognition in Experiment 2 was scored only for correctly recognised items, so source analyses were based on fewer trials than were presented.

##### Experiment 3

Scene memory accuracy was quantified using two measures: gist and identification, as detailed in the preregistration. The gist measure captured the main theme of the scene, including all elements that, if omitted, would change the semantic meaning of the scene. The identification measure assessed correct identification of a specific picture, distinguishing it from other scenes with similar elements within the stimulus set. Two independent raters scored scene memory descriptions for all participants. Interrater reliability was high for both measures and both cues: object cue: gist κ = 0.77, identification κ = 0.87; word cue: gist κ = 0.82, identification κ = 0.91.

Both gist and identification measures of recall were analysed in a mixed ANOVA with epoch condition (0, 2, Baseline) and cue type (word, object) as repeated measures factors. The amnesic shadow was defined as the difference in gist and identification between the two-epoch and baseline conditions, particularly when tested with the word cue. Because each scene was reactivated across both halves of the attention modulation phase, the data could neither be analysed across task halves nor tested for growth over time, unlike Experiments 1 and 2.

##### Online measures of divided attention

Online measures of attention were assessed using data from the odd/even buffer tasks and bystander tasks in the attention modulation phase. Following the amnesic shadow analysis approach, divided attention tests were conducted on data from the second half in Experiments 1 and 2, and across the whole task in Experiment 3. For the buffer tasks, reaction time and accuracy were computed depending on whether the digits followed a difficult or easy discrimination task. For bystander tasks, subjective judgements (story generation difficulty in Experiment 1) and an objective old/new recognition judgement (Experiment 3), along with their respective reaction times, were averaged and analysed across epoch conditions. In Experiment 2, only time to make animacy/pleasantness judgement ratings was analysed across epoch conditions because subjective ratings of pleasantness do not have a clear measure of accuracy.

##### General analysis approach

Statistical analyses were conducted in R (version 4.3.3) using RStudio (Cherry Blossom; v2023.3.0.386). Primary hypotheses were tested using ANOVAs. Significant omnibus tests were followed up with planned paired-samples t-tests where appropriate, corrected for multiple comparisons using Fisher’s least significant difference procedure. Item counterbalancing was included as a between-participants nuisance factor to account for variability associated with the stimulus materials in Experiments 1 and 3. Greenhouse–Geisser corrections were applied to all repeated measures effects with more than two levels. Effect sizes are reported as partial η*2* for ANOVAs and Cohen’s d for t-tests. Confidence intervals are reported for both F-tests (90%) and t-tests (95%)^114^. The alpha level was set at 0.05 (two-tailed). Bayes factor ANOVAs (default priors of the BayesFactor package, including a Cauchy prior scale of 0.5 for fixed effects, with at least 10,000 MCMC iterations) were used where evidence in favour of the null was of theoretical interest, specifically in the tests of divided attention.

#### Apparatus and Software

Experiment 1b and Experiment 3 used PsychoPy v2023.2.1 for stimulus presentation. Visual stimuli were presented on a 24-inch VPixx ViewPixx/3D monitor (1920 × 1080 pixels; 120Hz refresh rate), providing the temporal precision required for accurate stimulus timing in the brightness discrimination task. Experiments 1a and 2 used Presentation® software (Neurobehavioral Systems, Berkeley, CA) for stimulus presentation; hardware specifications for these setups are unavailable. Participants viewed the screen from a distance of 60cm across all experiments, maintained throughout the duration of the experiment.

#### Pre-registration

Experiments 1a and 2 were not preregistered. For dataset 1b, the sample size and scoring rules were preregistered as part of a replication attempt (Click here for link to preregistration); the analyses reported here pool datasets 1a and 1b and were therefore not preregistered. Experiment 3 was fully preregistered on OSF on 23rd October 2024 (Click [here](https://osf.io/k7rf5/overview?view_only=cdbf268ec2a84d96b28050e94f868661) for link to preregistration). There were no deviations from the preregistered plan.

#### Ethical Recruitment

The relevant ethics boards granted ethical approval for all experiments (Oregon: Institutional Review Board protocol #C1-314-07F; St. Andrews: University Teaching and Research Ethics Committee reference number PS5008; MRCCBU: Cambridge Psychology Research Ethics Committee reference number PRE.2023.068). Informed consent was obtained from all participants. Participants consented to the publication of anonymised data in peer-reviewed journals and its deposit in open repositories.

## Acknowledgements

We thank Napat Burapath and Gaia Austoni for their help with stimuli curation and data collection for Experiment 3. We thank all research assistants across University of Oregon, University of St. Andrews, and MRC Cognition and Brain Sciences Unit, University of Cambridge for their assistance with data collection. We thank Aleyna Namli for serving as the second rater for scene memory descriptions in Experiment 3. We also thank members of the Memory Control Lab at the MRC Cognition and Brain Sciences Unit for helpful discussions and feedback on this work. We thank Christian L. Ebbesen for the bioRxiv Word [template](https://github.com/chrelli/bioRxiv-word-template) used to format this preprint.

## Data availability

All data will be made available in a public repository after peer review.

## Code availability

All analysis scripts and the experiment task scripts will be made available in a public repository after peer review.

## Author contributions

A.S., J.C.H. and M.C.A. conceived and designed the study. J.C.H. and A.S. collected the data. A.S. analysed the data. A.S. and M.C.A. interpreted the results. A.S. and M.C.A. wrote the first draft of the manuscript, and J.C.H. reviewed and edited it. M.C.A. supervised the project and acquired funding. All authors approved the final version.

