## Supplementary Information for "Amnesia Induced by Suppressing the Focus of Internal Attention"

### Online tests of divided attention

#### *Analysis of buffer trial performance*

An alternative account for the observed memory deficits is that difficult discriminations divide attention greater than easy discriminations, and reduce processing of upcoming bystanders. Accordingly, divided attention predicts that difficult discriminations should differentially distract participants relative to easy discriminations. We analysed data across buffer trials following the visual discrimination task until bystander presentation (see Figure S1).

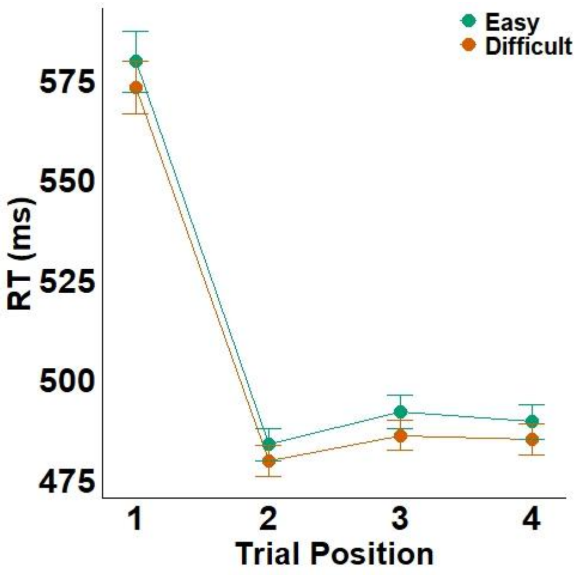


*Figure S1.* Reaction times across buffer-trial positions following difficult (orange) and easy (green) discriminations, leading up to bystander presentation, pooled across all three experiments. The task-switch cost from the discrimination task is evident in the markedly slower RTs at trial 1 relative to trials 2–4. At trial 1, reaction times were slightly faster following difficult than easy discriminations, but this difference was not evident across subsequent trials (2–4). Error bars represent standard error of the mean ($N$ = 194).

Analysis of buffer task data across all three experiments (second half of Experiments 1 and 2, and the whole task in Experiment 3; $N$ = 194) indicated that participants were not slower following difficult relative to easy discriminations, contrary to the predictions of divided attention.

First, we collapsed across the first four buffer trials following visual discriminations. Reaction times were reliably faster following difficult ($M$ = 503ms, $SD$ = 54.8) than easy ($M$ = 509ms, $SD$ = 61.3) discriminations ($F(1, 189) = 7.07$, $p = .009$, $\eta_p^2 = .036$, 90% CI = [.005, .09]), and buffer trial accuracy was reliably higher following difficult ($M$ = 92.9%, $SD$ = 11.2) than easy ($M$ = 91.5%, $SD$ = 12.5) discriminations ($F(1, 190) = 4.56$, $p = .034$, $\eta_p^2 = .023$, 90% CI = [.001, .07], $BF_{10} = 5.40$). We next verified that the buffer task was sensitive to the distracting effects of task-set switching. When reaction times were broken down by the serial position following discriminations, they were considerably longer for the first buffer trial that occurred immediately after a discrimination task ($M$ = 577ms, $SD$ = 99.4) compared to the remaining trials ($M$ = 483ms, $SD$ = 51.6; $t(189) = 21.16$, $p < .001$, $d = 1.11$, 95% CI = [0.93, 1.28]). Accuracy followed the same pattern, with the first buffer trial responded to less accurately ($M$ = 90.5%, $SD$ = 13.5) than the remainder ($M$ = 92.8%, $SD$ = 11.0; $t(190) = -2.96$, $p$ = .004, $d$ = −0.28, 95% CI = [−0.42, −0.13]). Hence, the buffer task was sensitive to detecting the task-switch cost following visual discriminations.

To scrutinise differential distraction, we examined performance on the very first buffer trial, which should be most sensitive to task-switches from the preceding discrimination. Divided attention would predict slower reaction times on this first trial following difficult relative to easy discriminations. We found evidence contrary to this prediction, as participants were reliably faster following difficult ($M$ = 573ms, $SD$ = 94) than easy ($M$ = 581ms, $SD$ = 112) discriminations ($F(1, 189) = 8.62$, $p$ = .004, $\eta_p^2$ = .044, 90% CI = [.008, .101], $BF_{10}$ = 0.77), and more accurate ($M$ = 91.2%, $SD$ = 13.6 vs. $M$ = 89.8%, $SD$ = 15.3; $F(1, 190) = 3.93$, $p$ = .049, $\eta_p^2$ = .020, 90% CI = [0.0001, .065], $BF_{10}$ = 0.75). The switch cost, operationally defined as the difference between the first and second buffer trial, was smaller following difficult ($M$ = 97ms, $SD$ = 82.7) than easy ($M$ = 103ms, $SD$ = 102) discriminations ($F(1, 189) = 3.97$, $p$ = .048, $\eta_p^2$ = .021, 90% CI = [0.0001, .066], $BF_{10}$ = 0.22). Reaction times across the remaining buffer trials (positions 2 to 4) did not reliably differ between difficult and easy discriminations ($M$ = 480ms, $SD$ = 51.2 vs. $M$ = 485ms, $SD$ = 58.0; $F(1, 189) = 2.93$, $p$ = .089, $\eta_p^2$ = .015, 90% CI = [0, .056], $BF_{10}$ = 0.64), nor did accuracy ($M$ = 93.5%, $SD$ = 11.3 vs. $M$ = 92.0%, $SD$ = 13.1; $F(1, 190) = 2.46$, $p$ = .119, $\eta_p^2$ = .013, 90% CI = [0, .052], $BF_{10}$ = 0.70).

Together, we confirmed that the buffer task was sensitive to the distracting effects of task switching, but we found no evidence that difficult discriminations differentially distracted participants relative to easy discriminations. Where reliable effects emerged, they indicated better performance following difficult relative to easy discriminations, the opposite of the predictions of divided attention.

#### *Analysis of bystander task performance*

Divided attention predicts worse performance in the bystander tasks themselves, following difficult relative to easy discriminations. We did not find evidence in support of this prediction in Experiment 1 (reported in the main text), nor in Experiments 2 or 3. In Experiment 2, reaction time to make subjective judgements did not differ across epoch conditions (epoch 0, $M$ = 757ms, SD = 174; epoch 1, $M$ = 831ms, $SD$ = 165; epoch 2, $M$ = 806ms, $SD$ = 217; $F(1.40, 39.23) = 2.85$, $p$ = .086, $\eta_p^2$ = .092, $BF_{10}$ = 0.66). In Experiment 3, recognition accuracy of old objects (bystanders) was similar between two and zero epoch conditions ($F(1,57) = 0.47$, $p$ = .496, $\eta_p^2$ = .008, $BF_{10}$ = 0.24), and so were reaction times ($F(1,57) = 1.30$, $p$ = .26, $\eta_p^2$ = .022, $BF_{10}$ = 0.30). Together, we do not find evidence that difficult discriminations reduced upcoming bystander processing relative to easy discriminations. Hence, these secondary analyses further did not provide evidence for divided attention.

#### **Buffer trials following bystander presentation**

The buffer trials completed after bystander tasks were intended to discourage participants from having lingering thoughts about the bystander, which could differentially improve subsequent memory performance. Such rehearsal following bystander tasks could be more likely after easy discriminations relative to difficult discriminations, improving bystander memory rather than impairing memory as we predicted.

However, we did not find evidence supporting this proposal. Collapsing across the first four after-bystander buffer trials (second half of Experiments 1 and 2, and the whole task in Experiment 3), reaction times did not reliably differ between conditions (Easy: *M* = 523ms, *SD* = 65.2; Difficult: *M* = 526ms, *SD* = 63.4; $F(1, 189) = 0.90$, $p = .345$, $\eta_p^2 = .005$, 90% CI = [0, .034], $BF_{10} = 0.33$), nor did accuracy (Easy: *M* = 91.0%, *SD* = 11.4; Difficult: *M* = 90.5%, *SD* = 11.7; $F(1, 190) = 0.93$, $p = .336$, $\eta_p^2 = .005$, 90% CI = [0, .035], $BF_{10} = 0.21$). Hence, there was no indication that participants differentially lingered on bystanders following easy relative to difficult discriminations.

Therefore, we did not find evidence that participants were slower in after-bystander buffer trials following easy discriminations, providing no evidence that participants were actively rehearsing bystander information during easy trials in a way that could benefit memory retention.

**Experiment 2 analyses**

We report relevant comparisons that did not yield statistically reliable results in Experiment 2. For source memory, comparisons between one and zero epochs ($F(1, 28) = 1.42$, $p = .243$, $\eta_p^2 = .048$, 90% CI = [0, .22]), or between two and one epochs ($F(1, 26) = 0.99$, $p = .330$, $\eta_p^2 = .037$, 90% CI = [0, .21]) did not reveal reliable differences.

**Experiment 3 analyses**

We found evidence consistent with the possibility that associative interference induced forgetting of bystander scenes reactivated with objects in the attention modulation phase relative to forgetting of baseline memories. When memory was tested with object cues, below-baseline forgetting was observed for bystanders in both the easy (gist $t(59) = 2.46$, $p = .017$, $d = .30$; identification $t(59) = 4.38$, $p < .001$, $d = .49$) and difficult conditions (gist $t(59) = 4.57$, $p < .001$, $d = .49$; identification $t(59) = 4.77$, $p < .001$, $d = .55$). We also did not find reliable differences between easy and difficult conditions specifically when tested with the object cues alone (gist $t(59) = 1.35$, $p = .184$, $d = .17$; identification $t(59) = 0.51$, $p = .614$, $d = .06$).
